# LaminaRGeneVis: a tool to visualize gene expression across the laminar architecture of the human neocortex

**DOI:** 10.1101/2021.05.12.443851

**Authors:** Ethan Kim, Derek Howard, Yuxiao Chen, Shreejoy J. Tripathy, Leon French

## Abstract

Application of RNA sequencing has enabled the characterization of genome-wide gene expression in the human brain, including distinct layers of the neocortex. Neuroanatomically, the molecular patterns that underlie the laminar organization of the neocortex can help link structure to circuitry and function. To advance our understanding of cortical architecture, we created *LaminaRGeneVis*, a web application that displays across-layer cortical gene expression from multiple datasets. These datasets were collected using bulk, single-nucleus, and spatial RNA sequencing methodologies and these data were harmonized to facilitate comparisons between datasets. The online resource facilitates single- and multi-gene analyses by providing figures and statistics for user-friendly assessment of laminar gene expression patterns in the adult human neocortex.

**Availability and implementation:** *LaminaRGeneVis* is available at https://ethanhkim.shinyapps.io/laminargenevis. The source code and data is accessible at https://github.com/ethanhkim/laminargenevis.

## 1. Introduction

RNA sequencing has provided molecular markers of human brain anatomy by revealing spatial gene expression patterns. The application of these techniques has provided genome-wide profiles of expression, but the brain’s complexity has limited our understanding of its cellular, molecular, and laminar architecture. Specifically, while the cytoarchitecture of the recently evolved neocortex has been characterized, we lack a strong understanding of its layer-specific gene expression.

Currently, few web applications visualize gene expression in the adult human neocortex. The Allen Brain Atlases from the Allen Institute of Brain Science (AIBS) and other tools allow viewing spatial expression patterns (Guo et al., 2019; Maynard et al., 2021; Shen et al., 2012; Zeng et al., 2012). However, there are no visualization tools to analyze expression across human neocortical layers. Several datasets provide this laminar data but due to differences in the scopes and methods used, accessing and comparing this data is difficult and time-consuming.

Here, we present LaminaRGeneVis, a web application for analyses of gene expression across human neocortical layers. LaminaRGeneVis enables visualization and analysis of data from layer-specific bulk-tissue, single-nucleus, and spatial transcriptomic RNA sequencing studies.

## 2. Data and Methods

### 2.1 Datasets

We used data from three studies that assayed genome-wide expression across the layers of the human neocortex in neurotypical donors. First, He and colleagues transversely sliced dorsolateral prefrontal cortex samples (DLPFC) from postmortem brains (He et al., 2017). Guided by their analyses, we focused on their first dataset (DS1) which contains expression data from four brains and was obtained from SRA using project code SRP065273. A second study of the DLPFC from 3 adult donors employing the spatial transcriptomics 10X Genomics Visium platform was obtained from the spatialLIBD R package (Maynard et al., 2021). The third dataset is from AIBS and assayed expression with single-nucleus RNA sequencing (snRNA-seq) in 3 brains (https://portal.brain-map.org/atlases-and-data/rnaseq/human-multiple-cortical-areas-smart-seq). We chose to use data only from the middle temporal gyrus, as this region had the most samples. The data was also split into three cell types as labelled by AIBS: GABAergic, glutamatergic and non-neuronal. While methods for spatial dissection vary, all three of these studies employed RNA sequencing and profiled the adult human neocortex. The characteristics of these datasets are described in Supplementary Table 1. Similarity analyses were performed on the datasets to validate subsequent expression correlation and layer-specific enrichment analyses. The results of those analyses are available in Supplementary Results, Tables S2 and S3.

#### 2.1.1 Dataset processing

Each dataset was processed using a similar pipeline as described in Supplementary Methods. This processing resulted in each dataset being represented in a gene expression matrix in the shape of genes (rows) by layer (columns), with gene expression represented as normalized and log-transformed read counts (log2(counts per million)).

### 2.2 Statistical analysis

#### 2.2.1 Gene expression correlation

Pearson correlation was used to assess agreement for single genes in the bulk-tissue data. This is calculated using a given gene’s seven layer-specific expression values from the He and Maynard datasets. For multiple genes, average correlation is used. To add a genome-wide perspective, the percentile of these correlations in reference to all other gene-to-gene values are also calculated.

#### 2.2.2 Layer-specific gene set enrichment

To test for the enrichment of a set of genes in a given layer, we first removed genes with CPM < 0.1 across all layers, and normalized through log2-transformation then z-score normalized. We sorted the normalized expression matrices per layer by ranking the remaining genes in each dataset from the most to least normalized expression. Within this ranking, the area under the receiver operating characteristic curve (AUC) was used to test whether the inputted set of genes were enriched or depleted (enrichment: AUC > 0.5, depletion: AUC < 0.5). The Mann–Whitney U test was used to determine statistical significance and multiple-test corrected with Bonferroni correction.

#### 2.2.3 Availability of data and code

Data used in the application and the code to process the data are available online at https://github.com/ethanhkim/laminargenevis. Scripts to process the raw data from He et al. are available at https://github.com/derekhoward/he_seq.

### The LaminaRGeneVis application

*LaminaRGeneVis* is a web application implemented in shiny (Chang et al., 2021) that can be accessed at https://ethanhkim/shinyapps.io/laminargenevis. Users can choose either Single or Multi-gene mode to visualize gene expression for one or a set of genes, respectively. The application will also report statistical analyses that assay agreement in the bulk-tissue data and test layer-specific enrichment for a set of genes.

#### Single gene mode

The user can input their gene of interest by typing in its gene symbol and selecting it from the drop-down bar. Once submitted, the gene’s normalized expression across the cortical layers in each dataset is displayed as a bar plot. A text box below notes which datasets assayed the queried gene and the agreement statistics in bulk-tissue datasets.

#### Multi-gene mode

In the Multi-gene mode, the application generates visualizations for the queried gene set’s layer-specific enrichment and normalized expression of each gene across datasets.

Layer-specific enrichment is visualized as a heatmap that displays the AUC values across layers and datasets. Normalized gene expression is displayed as a heatmap for 30 genes or less and a scatterplot otherwise. The scatterplots also show the queried genes’ median expression in each layer. A summary textbox at the bottom reports information such as the number of inputted genes assayed in each bulk-tissue dataset and dataset agreement statistics across the bulk-tissue data.

## Conclusion

We have developed a web application for visualizing gene expression across the laminar architecture of the adult human neocortex. It reports cross-dataset correlation and the enrichment of layer-specific expression. These functionalities provide easily accessible figures and statistics for quick assessment of expression across the layers of the human cortex.

## Acknowledgements

The work described was supported by funding from the Natural Sciences and Engineering Research Council of Canada and Brain Canada, in partnership with Health Canada, for the Canadian Open Neuroscience Platform.

---

**Fig. 1.**
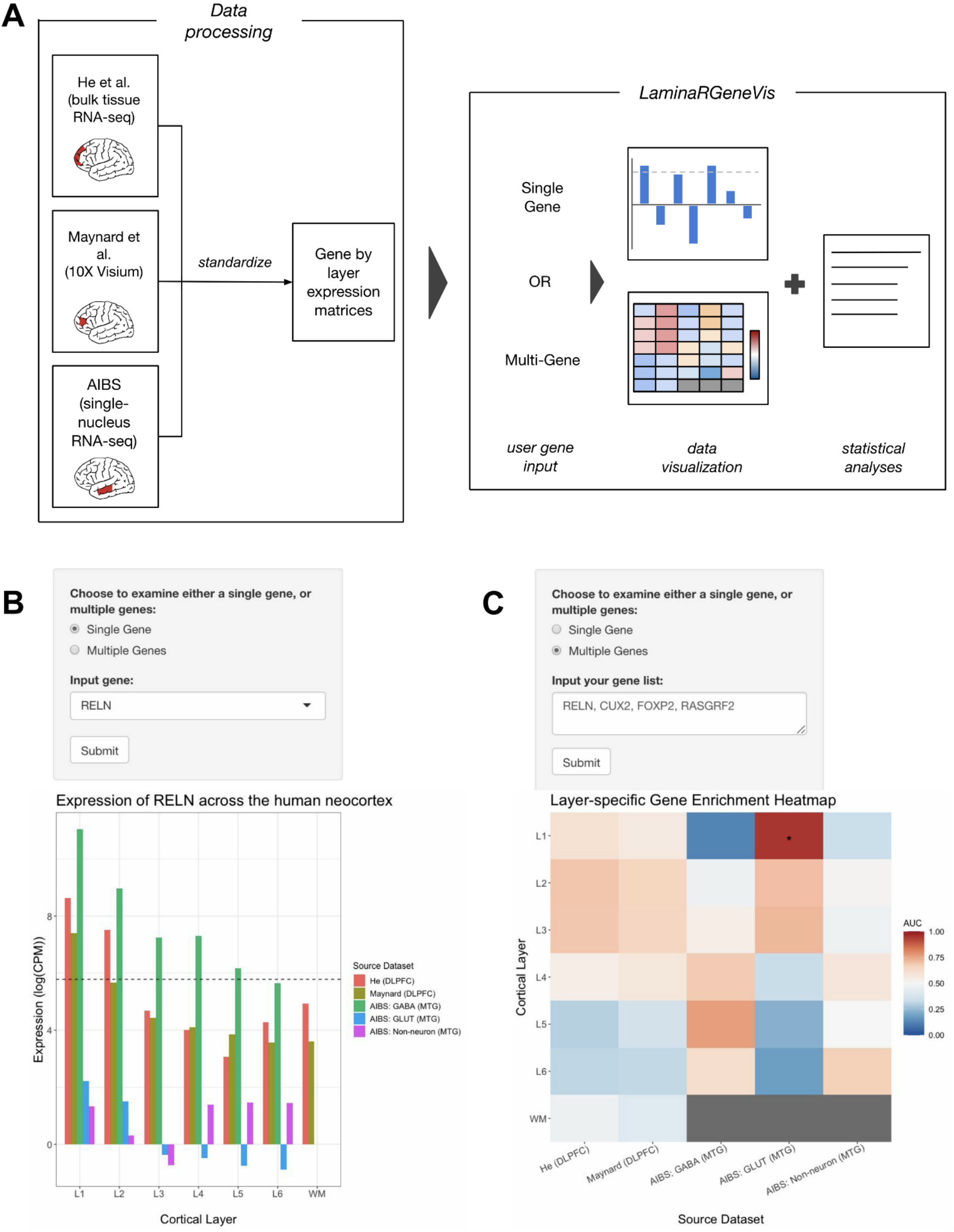
Workflow diagram and visualizations of gene expression across the cortical layers. (**A**) Diagram of data flow and application usage. Each dataset was uniformly processed and standardized to create normalized gene expression matrices. These matrices are used in the web application for visualizations and statistical analyses. Dataset characteristics are summarized in Supplementary Table S1. **(B)** When the user chooses to examine a single gene with the selection options, the application will display a bar plot showing the normalized expression of the gene across the cortical layers and white matter. **(C)** Upon choosing to examine multiple genes, the application will visualize the genes’ layer-specific enrichment (as shown) as a heatmap with asterisks marking statistically significant AUC scores.

