## Supplementary Material for "LaminaRGeneVis: a tool to visualize gene expression across the laminar architecture of the human neocortex"

*Supplementary Methods*

Dataset processing

The raw RNA-seq data from He et al. was downloaded from Sequence Read Archive (project code SRP065273) in .fastq format. Genome alignment was performed using STAR 2.7 (Dobin et al., 2013) against the reference genome GRCh38.p13 with the corresponding annotation file from Ensembl (Yates et al., 2020). Default parameters were used. Aligned reads were quantified using RSEM (Li & Dewey, 2011) against the same reference genome and annotation file. This created a count matrix of 59,453 genes across 102 samples. We focused our analyses on selected samples from Dataset 1 labelled (DS1) to mirror the focus of He et al. In instances where multiple Ensembl ID's referred to the same HGNC gene symbol, we took the average of the counts. For each of the samples that were labelled as the same transverse slice (e.g., all samples with the S1 label for slice 1), we summed the counts across the sample columns for all 18 transverse slices. The resulting 18 columns were pseudo-bulked at the layer level to create 7 columns representative of the 6 layers of the human neocortex and 1 white matter layer using the mapping provided by He and colleagues (weighted averages). To avoid taking the log of 0, we added 1 to all counts prior to counts per million (CPM) normalization. We used the cpm() function from the edgeR package (Robinson et al., 2010; McCarthy et al., 2012) to CPM normalize the data, using log = T to $\log_2$ transform the data. The final normalized expression matrix contains 59,453 genes' expression in the 6 neocortical layers and 1 layer of white matter in the DLPFC.

Maynard et al. created an R package, spatialLIBD, which allows for researchers to download the raw data from their use of the Visium platform. We used the fetch_data() function in their package with type = “sce_layer” to download the sce_layer data, which is the pseudo-bulked expression matrix containing gene expression data for 22,331 genes across 12 samples spanning the 6 neocortical layers and 1 white matter layer. We only used the samples from two of the three donors (sample ID’s: 151507, 151508, 151509, 151510, 151673, 151674, 151675, 151676) that contained data for all 6 cortical layers, reducing the number of usable samples to 8. We used a similar aggregation and normalization pipeline used for the He et al data, but altered the summation across samples such that we took the sum across samples labeled as the same cortical layer. The final matrix contains 18,633 genes’ expression in the 6 neocortical layers in the DLPFC, as well as a layer of white matter similar to the He et al. matrix.

The Allen Brain Cell Types database contains multiple single nucleus RNA-seq datasets, from which we chose to use the “Multiple Cortical Area - SMART-seq (2019)” dataset, accessed from https://portal.brain-map.org/atlases-and-data/rnaseq/human-multiple-cortical-areas-smart-seq. This dataset contains data from 3 donors with multiple cortical regions assayed. We chose to only use the data from the middle temporal gyrus, as this region contained the highest number of nuclei (n = 15,519) and was the most similar to the layer labeling found in He et al and Maynard et al. The count matrix provided nucleus-level data, with cell-type and layer labels per nucleus. Using the provided metadata, we first filtered out samples with outlier_call = TRUE to remove any outlier nuclei. We then randomly downsampled the data for each of the cell types such that each layer contained the same number of nuclei per cell type. This resulted in 381, 274 and 125 nuclei per layer in the GABAergic, glutamatergic and non-neuronal data, respectively. The count matrix was pseudo-bulked at the level of layers by summing the raw gene-expression counts across all samples for each layer, cell-type and gene. The bulked dataset was normalized through CPM normalization and $log_2$-transformation, similar to the He and Maynard datasets. We then separated the dataset by cell-type to create three distinct normalized expression matrices of 50,286 genes across the 6 neocortical layers of the MTG.

*Supplementary Results*

Dataset Similarity

We evaluated dataset similarity using the Mantel test (Mantel, 1967). We used the implementation provided by the vegan R package (Oksanen et al., 2020), using Pearson correlation as the statistical method. As the Mantel test correlates similarity or distance matrices, We used gene-gene Pearson correlation matrices for the similarity matrices to test if across layer gene associations are consistent within and across the datasets.

We first examined the inter-and intra-donor similarity in the Maynard data for the two donors with all 6 cortical layers and white matter layer labelled. We downloaded the layer-level data as previously described, from which we separated each sample from the two donors and normalized through CPM normalization and log2-transformation.

This resulted in 4 expression matrices per donor for a total of 8 matrices. The mean Mantel r statistic for donor 1 was 0.4106 and 0.4140 for donor 2. When comparing across donors, the mean r statistic was 0.2947. We validated these results by subsetting the Maynard data to a list of genes used by Zeng et al (Zeng et al., 2012). This resulted in a mean r statistic of 0.5303 for donor 1 and 0.6456 for donor 2. The mean r statistic across donors was 0.4324. All sample-to-sample comparisons can be found in Supplementary Table 2.

We then compared the similarity between the Maynard and He datasets using the bulked and normalized datasets as described in Supplementary Methods, Dataset processing section. We subsetted the datasets to a list of genes common between the two (n = 18,206), and each dataset was correlated to itself to create correlation matrices for Mantel testing. We found that the Mantel r statistic was 0.5166 when comparing across datasets. Removing genes with CPM < 0.1 prior to log-transformation resulted in the r statistic increasing to 0.5731 (15,589 genes). We improved these results by subsetting the two datasets to a list of layer-specific marker genes (Zeng et al., 2012). We found that the r statistic was 0.683 (946 genes) and 0.7246 when filtering for genes with CPM > 0.1 (882 genes).

We also compared the bulk-tissue datasets to the cell-type specific snRNA-seq data from the AIBS using the aforementioned processes for comparing the Maynard and He data. We found after removing white matter data from the bulk-tissue, there was little to no correlation in either direction between the bulk-tissue and cell-type specific snRNA-seq data: Comparing the He dataset against each of the cell-type specific data resulted in a mean r statistic of -0.08134 (30,744 genes) and 0.0007667 for Maynard et al. (17,197 genes). When filtering for genes with CPM > 0.1 across all layers, there were slight increases in the r statistic and subsetting to the list of genes from Zeng et al. also slightly increased the r statistic.

*Supplementary Table 1: Characteristics of the datasets used in LaminaRGeneVis*

| **Dataset** | **Technique used** | **Type of samples used:** | **Gene expression quantificatio n** | **Cortical region assayed:** | **# of genes assayed** | **# of donors** |
|---|---|---|---|---|---|---|
| He et al. | Illumina RNA-seq | Bulk-tissue | Gene count | DLPFC (BA 9, 10) | 59,453 | 4 M |
| Maynard et al. | Visium (10X Genomics) | Tissue sections | Raw UMI count | DLPFC (BA 46) | 18,633 | 2 (1 M, 1 F) |
| Allen Cell Type Database | SMART-seq RNA-seq | Single nuclei | Gene count | MTG | 50,286 | 3 (2 M, 1 F) |

** DLPFC, dorsolateral prefrontal cortex; MTG, middle temporal gyrus; BA, Brodmann Area*

*Supplementary Table 2: Dataset similarity testing results for Maynard et al data*

| **Donor label 1** | **Donor label 2** | **Sample IDs tested** | **Mantel statistic r** | **Mantel statistic r; filtered for Zeng et al marker list** |
|---|---|---|---|---|
| 1 | 1 | 151507, 151508 | 0.3867437 | 0.4929938 |
| 1 | 1 | 151507, 151509 | 0.3761705 | 0.4928974 |
| 1 | 1 | 151507, 151510 | 0.3662345 | 0.5029497 |
| 1 | 1 | 151508, 151509 | 0.4472683 | 0.5596710 |
| 1 | 1 | 151508, 151510 | 0.4217159 | 0.5252885 |
| 1 | 1 | 151509, 151510 | 0.4652534 | 0.6079797 |

| | | | | |
|---|---|---|---|---|
| 2 | 2 | 151673, 151674 | 0.4450776 | 0.6862513 |
| 2 | 2 | 151673, 151675 | 0.4396620 | 0.6846133 |
| 2 | 2 | 151673, 151676 | 0.4170934 | 0.6318101 |
| 2 | 2 | 151674, 151675 | 0.4021281 | 0.6449719 |
| 2 | 2 | 151674, 151676 | 0.3875977 | 0.6121642 |
| 2 | 2 | 151675, 151676 | 0.3922045 | 0.6138546 |
| 1 | 2 | 151507, 151674 | 0.3079286 | 0.4083163 |
| 1 | 2 | 151507, 151673 | 0.2747824 | 0.4145270 |
| 1 | 2 | 151507, 151675 | 0.2853121 | 0.4195137 |
| 1 | 2 | 151507, 151676 | 0.2729248 | 0.4080120 |
| 1 | 2 | 151508, 151674 | 0.3110389 | 0.3904035 |
| 1 | 2 | 151508, 151673 | 0.2602919 | 0.3850534 |
| 1 | 2 | 151508, 151675 | 0.2833523 | 0.3833158 |
| 1 | 2 | 151508, 151676 | 0.2703848 | 0.3671039 |
| 1 | 2 | 151509, 151674 | 0.3625330 | 0.5043142 |
| 1 | 2 | 151509, 151673 | 0.3021198 | 0.4848182 |
| 1 | 2 | 151509, 151675 | 0.3207013 | 0.4837968 |
| 1 | 2 | 151509, 151676 | 0.3041808 | 0.4514819 |
| 1 | 2 | 151510, 151674 | 0.3313964 | 0.4796648 |
| 1 | 2 | 151510, 151673 | 0.2823863 | 0.4555763 |
| 1 | 2 | 151510, 151675 | 0.3066032 | 0.4513501 |
| 1 | 2 | 151510, 151676 | 0.2389498 | 0.4309212 |

*Supplementary Table 3: Dataset similarity testing results for Maynard et al, He et al and Allen Institute data*

| **Dataset 1** | **Dataset 2** | **Mantel statistic r** | **Mantel statistic r; filtered for CPM > 0.1** | **Mantel statistic r; filtered for Zeng et al list** |
|---|---|---|---|---|

| | | | | |
|---|---|---|---|---|
| He et al | ACTD - GABAergic | -0.1077 | -0.006967 | 0.03838 |
| He et al | ACTD - Glutamatergic | -0.1295 | -0.005718 | 0.03376 |
| He et al | ACTD - Non-neuronal | -0.006967 | 0.05965 | 0.07922 |
| He et al | ACTD - layer-aggregate | 0.03536 | 0.09435 | 0.1756 |
| Maynard et al | ACTD - GABAergic | -0.01428 | 0.002822 | 0.03121 |
| Maynard et al | ACTD - Glutamatergic | -0.0143 | 0.009769 | 0.064 |
| Maynard et al | ACTD - Non-neuronal | 0.02628 | 0.04014 | 0.06274 |
| Maynard et al | ACTD - layer-aggregate | 0.08748 | 0.1037 | 0.201 |
| He et al | Maynard et al | 0.5166 | 0.5731 | 0.683 |

** ACTD Allen Cell Type Database; MTG, middle temporal gyrus; BA, Brodmann Area*
